# A dual-metric framework for quantifying the biological fidelity of scRNA-seq pipelines

**DOI:** 10.1101/2025.11.12.688141

**Authors:** Qiang Su, Yi Long, Fuyu Duan, Qizhou Lian

**Affiliations:** Faculty of Synthetic Biology, Shenzhen University of Advanced Technology, Shenzhen 518107, China; Faculty of Life and Health Sciences, Shenzhen University of Advanced Technology, Shenzhen 518107, China; State Key Laboratory of Quantitative Synthetic Biology, Shenzhen Institute of Synthetic Biology, Shenzhen Institutes of Advanced Technology, Chinese Academy of Sciences, Shenzhen 518055, China

**Keywords:** scRNA-seq, Biological fidelity, Computational Pipeline, CAS-MCS-Scoring

## Abstract

Single-cell RNA sequencing (scRNA-seq) pipelines differ in barcode recovery, UMI correction, and read assignment, potentially affecting downstream biological interpretation. We developed a reproducible, parameter-aware framework to benchmark STARsolo, Cell Ranger, Kallisto|Bustools, and Simpleaf using simulated datasets and biological samples, including a Huntington’s disease mouse model. We introduced two complementary metrics: the Cluster Annotation Score (CAS), which measures agreement between direct cell-level and cluster-level annotations, and the Marker Concordance Score (MCS), which evaluates marker-gene coherence within cell types. By varying the numbers of highly variable genes and principal components, we assessed interactions between primary processing and downstream analysis. STARsolo and Kallisto|Bustools showed high technical accuracy in simulations, but STARsolo most consistently preserved stable cell identities and marker coherence. Cell Ranger also performed strongly, whereas Simpleaf showed variable performance across cell types and analytical settings. Kallisto|Bustools generally showed lower biological concordance. Our open-source toolkit supports transparent and biologically informed pipeline evaluation.

## Introduction

The challenge of unraveling cellular transcriptomes within heterogeneous tissues is fundamental to advancing our understanding of development, physiology, and disease.(Tirosh et al. 2016; Venteicher et al. 2017; Papalexi and Satija 2018; Van de Sande et al. 2023) Single-cell RNA sequencing (scRNA⍰seq) and single-nucleus RNA sequencing (snRNA⍰seq) represent transformative solutions, profiling millions of individual cells to illuminate heterogeneity and identify rare cell states missed by bulk analyses.(Klein et al. 2015; Macosko et al. 2015; Gawad et al. 2016; Lake et al. 2016; Habib et al. 2017; Zheng et al. 2017; Ziegenhain et al. 2017; Svensson et al. 2018; Wolf et al. 2018) These technologies derive their power from molecular barcoding, which uses cell barcodes (CBs) to denote cell of origin and unique molecular identifiers (UMIs)(Islam et al. 2014) to quantify individual transcripts.(Kivioja et al. 2011; Kolodziejczyk et al. 2015; Rosenberg et al. 2018; Ding et al. 2020) While barcoding enables high-throughput sequencing, the resulting gene-by-cell count matrix depends on computational processing of the raw reads. (Hwang et al. 2018; Gao et al. 2021) Because this matrix is the input to downstream analyses, differences introduced during primary processing can propagate to clustering, cell-type annotation, and biological interpretation.

The current ecosystem of primary processing pipelines is dominated by two distinct philosophies.(You et al. 2021; Pool et al. 2023) The first, exemplified by the industry-standard 10x Genomics Cell Ranger(Ding et al. 2020) and its open-source counterpart STARsolo,(Dobin et al. 2013; Kaminow et al. 2021; Moiseeva et al. 2023) relies on high-precision spliced genomic alignment. While computationally intensive, this approach provides granular, base-level resolution. These tools differ, however, in their statistical models for correcting barcode errors and collapsing UMIs-choices that directly impact cell recovery and quantification accuracy. The second philosophy prioritizes computational efficiency, using ultra-fast pseudoalignment (Kallisto|Bustools)(Melsted et al. 2021; Michelson et al. 2022; Weatherbee et al. 2023) or quasi-mapping via Simpleaf (a unified framework that standardizes and streamlines the execution of the Alevin-fry ecosystem)(He et al. 2022; He and Patro 2023; Gayoso et al. 2024) to rapidly assign reads to transcripts. This speed comes with its own trade-offs, as these methods employ more conservative, and algorithmically distinct, strategies for barcode and UMI error correction. Despite their widespread adoption, there is no consensus on how these algorithmic differences translate to biological reliability.(Slovin et al. 2021) This uncertainty is compounded by downstream analytical flexibility. Key parameters, such as the number of highly variable genes or principal components used for dimensionality reduction, are themselves subject to user choice. Crucially, it remains unknown how these downstream parameters interact with the outputs of different upstream pipelines, creating a complex analytical space where the path from raw reads to biological insight is opaque and non-standardized.

In this study, we systematically evaluated Cell Ranger, STARsolo, Kallisto|Bustools, and Simpleaf within a unified, parameter-aware analytical framework. Our aim was to move beyond conventional technical benchmarking and distinguish computational accuracy from downstream biological fidelity. We first assessed technical performance using a ground-truth simulated dataset, including barcode recovery and quantification accuracy. We then evaluated biological performance using complex experimental datasets, including public 10x Genomics resources and a Huntington’s disease mouse model, while systematically varying n_HVGs and n_PCs. This design allowed us to examine how upstream quantification interacts with downstream analytical parameters. To quantify complementary aspects of biological fidelity, we implemented two metrics in the open-source CAS-MCS-Scoring.py toolkit. The Cluster Annotation Score (CAS) measures concordance between direct cell-level annotations and cluster-level consensus annotations. The Marker Concordance Score (MCS) evaluates the coherence of annotated cell types by measuring the prevalence of de novo marker genes using a Wilcoxon rank-sum test (Gulati et al. 2025). Together, these metrics provide complementary measures of annotation concordance and marker-gene coherence, rather than serving as independent ground truth for cell identity. Our results distinguish technical performance from downstream biological concordance. Although STARsolo and Kallisto|Bustools performed strongly in the simulated technical benchmarks, their performance differed in empirical measures of annotation stability and marker coherence. Across the evaluated datasets, STARsolo and Cell Ranger generally showed strong annotation concordance, whereas Kallisto|Bustools frequently exhibited lower concordance. Simpleaf showed high CAS for multiple cell types and parameter settings, but its performance varied across cell populations and analytical conditions. These findings demonstrate that technical accuracy alone does not fully predict biological fidelity and that both primary-processing choice and downstream parameter selection contribute to the reproducibility and interpretation of scRNA-seq and snRNA-seq analyses.

## Results

### Algorithmic divergence across primary scRNA-seq processing pipelines

Analysis of scRNA-seq data follows a canonical workflow, yet the initial conversion of raw reads to a gene-by-cell count matrix is a critical juncture with multiple competing methodologies. To understand and quantify the impact of these choices on biological interpretation, we first dissected the algorithmic strategies of four widely adopted pipelines: Cell Ranger, STARsolo, KBus, and Simpleaf (Fig. 1a). Notably, in this study we utilized Simpleaf in place of standalone Alevin-fry; Simpleaf serves as a modern, unified framework that streamlines the execution of the Alevin-fry ecosystem, ensuring optimal, highly reproducible parameter usage while reflecting current best practices for lightweight mapping. While these tools share common goals, they diverge fundamentally in their approaches to read mapping, cell barcode demultiplexing, and UMI deduplication, representing distinct philosophies with significant theoretical trade-offs.

**Fig. 1.**
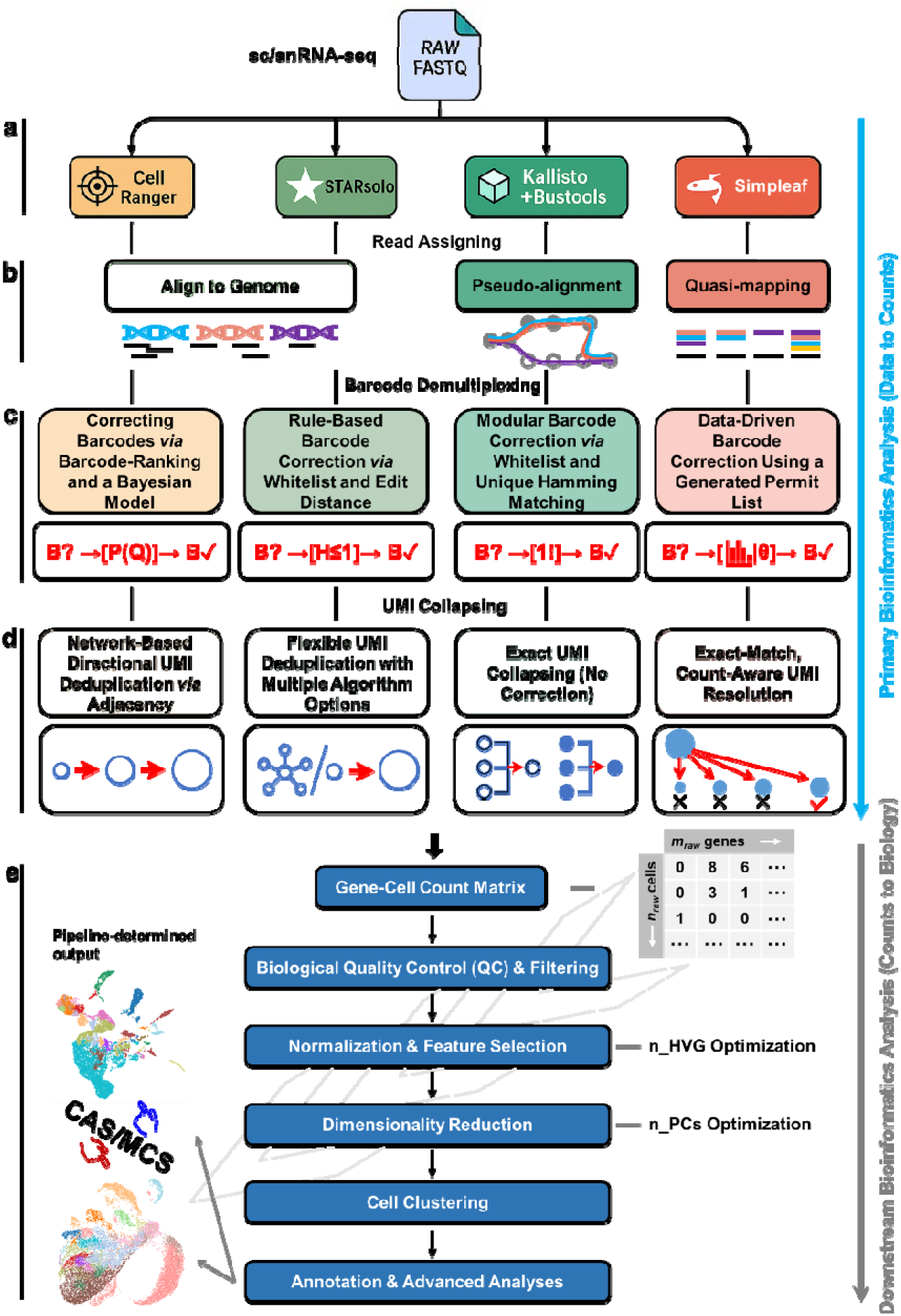
The canonical scRNA-seq analysis pipeline and its critical decision points. The workflow from raw data to biological interpretation highlights key variations at each stage. **a**) The process typically begins with one of four primary toolsets: Cell Ranger, STARsolo, KBus, and Simpleaf. **b**) These tools first differ in their core read processing strategy, with Cell Ranger and STARsolo using alignment-based methods while KBus and Simpleaf adopt alignment-free approaches. **c**) Next, each tool applies a distinct algorithm for cell barcode correction: Cell Ranger uses a Bayesian model, STARsolo employs rule-based edit distance, KBus performs unique Hamming matching, and Simpleaf generates a data-driven permit list. **d**) The subsequent UMI deduplication step is also tool-specific, involving network-based directional collapsing for Cell Ranger, flexible algorithm options for STARsolo, adjacency graph collapsing for KBus, and probabilistic resolution for Simpleaf. **e**) This primary analysis yields a gene-by-cell matrix that then undergoes a downstream workflow (e.g., QC, feature selection, data-dimension reduction, clustering), which itself contains tunable parameters. Consequently, the final biological interpretation is a pipeline-determined output, shaped by the combination of the chosen primary toolset and downstream parameter settings. This study aims to comprehensively benchmark these configurations to identify the pipelines providing the most accurate biological representation, evaluated using a dual-metric framework: the CAS for internal annotation stability and the MCS for gene-level biological validation.

The first major divergence occurs at the level of read processing and gene assignment (Fig. 1b). Cell Ranger and STARsolo use a spliced genomic alignment strategy, mapping each read with base-level precision to a reference genome via the STAR aligner. In contrast, KBus and Simpleaf adopt substantially faster k-mer-based methods-pseudoalignment and quasi-mapping, respectively-which bypass nucleotide-by-nucleotide alignment to identify transcript compatibilities. This fundamental architectural choice, trading mapping granularity for computational efficiency, directly influences not only performance but also the ultimate gene-level counts assigned to each cell, a key source of variability we sought to quantify.

Another critical stage is cell barcode demultiplexing (Fig. 1c), which identifies reads from true cells and corrects for sequencing errors. The pipelines employ distinct computational strategies: probabilistic inference in Cell Ranger; deterministic edit-distance matching in STARsolo; uniqueness-constrained correction in KBus; and adaptive, data-driven thresholding in Simpleaf. The chosen method directly influences the sensitivity and specificity of cell recovery, determining the final set of barcodes passed to downstream analysis. The final major stage is UMI deduplication (Fig. 1d), which corrects for amplification bias by collapsing reads from the same original RNA molecule. Here, strategies range from active, sequence-based error correction using abundance-aware graph approaches (Cell Ranger) or heuristics (STARsolo) to no explicit sequence correction (KBus, Simpleaf). These different approaches to handling UMI errors reflect distinct trade-offs between removing technical noise and the risk of over-collapsing true biological molecules, ultimately shaping the quantitative accuracy of the final count matrix. Full methodological details for all three stages are provided in the Methods section.

To systematically evaluate how these algorithmic divergences affect biological interpretation, we designed a comprehensive empirical evaluation framework. First, we processed the count matrix from each pipeline through a standardized downstream workflow (Fig. 1e), allowing us to benchmark foundational cell annotation accuracy, per-cell marker gene assignment, and internal cluster consistency. Second, we applied this framework to diverse empirical data to investigate the interplay between upstream pipeline choice and key downstream parameters-specifically the number of highly variable genes (n_HVG) and principal components (n_PCs). This approach enabled us to deconvolve the effects of initial quantification from downstream analysis, revealing how their interaction shapes everything from cluster resolution to final cell type annotation.

### How primary processing and downstream analysis decisions shape cluster-level cell type annotation in empirical data

An end-to-end scRNA-seq analysis framework transforms raw sequencing data into a biologically annotated single-cell atlas through a series of sequential steps (Fig.□2a). The process begins with primary data processing, where raw reads are quantified into a gene-by-cell count matrix by one of several pipeline-specific methods (e.g., STARsolo, Cell Ranger, KBus, or Simpleaf). This initial step is critical: methodological differences at this stage can propagate through all subsequent analyses, potentially altering the final biological interpretation. The downstream workflow then applies key tunable parameters-most notably n_HVG and n_PCs-to drive dimensionality reduction, clustering, and annotation. Cells are first grouped into clusters using the Leiden algorithm, after which each cluster is assigned a consensus cell type via a “majority vote” across individual cell-level predictions.

To systematically determine the optimal parameter settings for clustering in the downstream pipeline, we performed a simultaneous grid search over two key variables in the downstream analysis: n_HVG and n_PCs. For each primary processing method, we computed the Silhouette Score-an unsupervised metric of cluster separation quality-across a range of parameter combinations using the same 10x Genomics sourced 5k Adult Mouse Brain dataset. Across all primary processing methods, the resulting Silhouette Score landscapes exhibited method-specific plateau patterns (Fig.□2b). A common trend across all methods was a gradual decline in clustering quality as the number of PCs increased, regardless of the n_HVG setting. However, the optimal n_HVG value and the corresponding clustering topology were highly method-dependent. These findings highlight the critical interplay between primary processing methods and downstream parameter optimization, underscoring the need for method-specific tuning to achieve robust cluster resolution.

To investigate how downstream parameter choices affect clustering, we applied a highly restrictive setting (n_HVG = 51, n_PCs = 30) to gene-by-cell matrices from the four primary pipelines using the 10x Genomics 5k Adult Mouse Brain dataset (Fig. 2c). A low n_HVG setting disproportionately selects genes with high variance, which in scRNA-seq often correspond to a single dominant biological process-such as the cell cycle-rather than stable, cell-type-specific markers (Buettner et al. 2015). Consequently, clustering based on these few genes is expected to segment cells along a continuous biological trajectory rather than separating them into discrete cell types. This trajectory-driven artifact was acutely evident in the results from KBus, which collapsed the dataset into merely 8 broad clusters with exceptionally poor internal consistency (e.g., a massive 4,030-cell “Astro-TE” cluster with a CAS of just 14.81%). In stark contrast, STARsolo, Cell Ranger, and Simpleaf were much more resilient to this restricted feature space. They yielded lower, more realistic Silhouette Scores (0.106, 0.028, and 0.097, respectively), indicating that their underlying data structures were less dominated by trajectory-related variance. Despite the suboptimal conditions, these three pipelines successfully resolved a comparable and higher number of distinct cell types (16, 14, and 15, respectively) while maintaining moderate to high CAS scores across major populations (e.g., 72.14% for STR D1 Gaba in STARsolo; 74.47% for Oligo NN in Simpleaf).

**Fig. 2.**
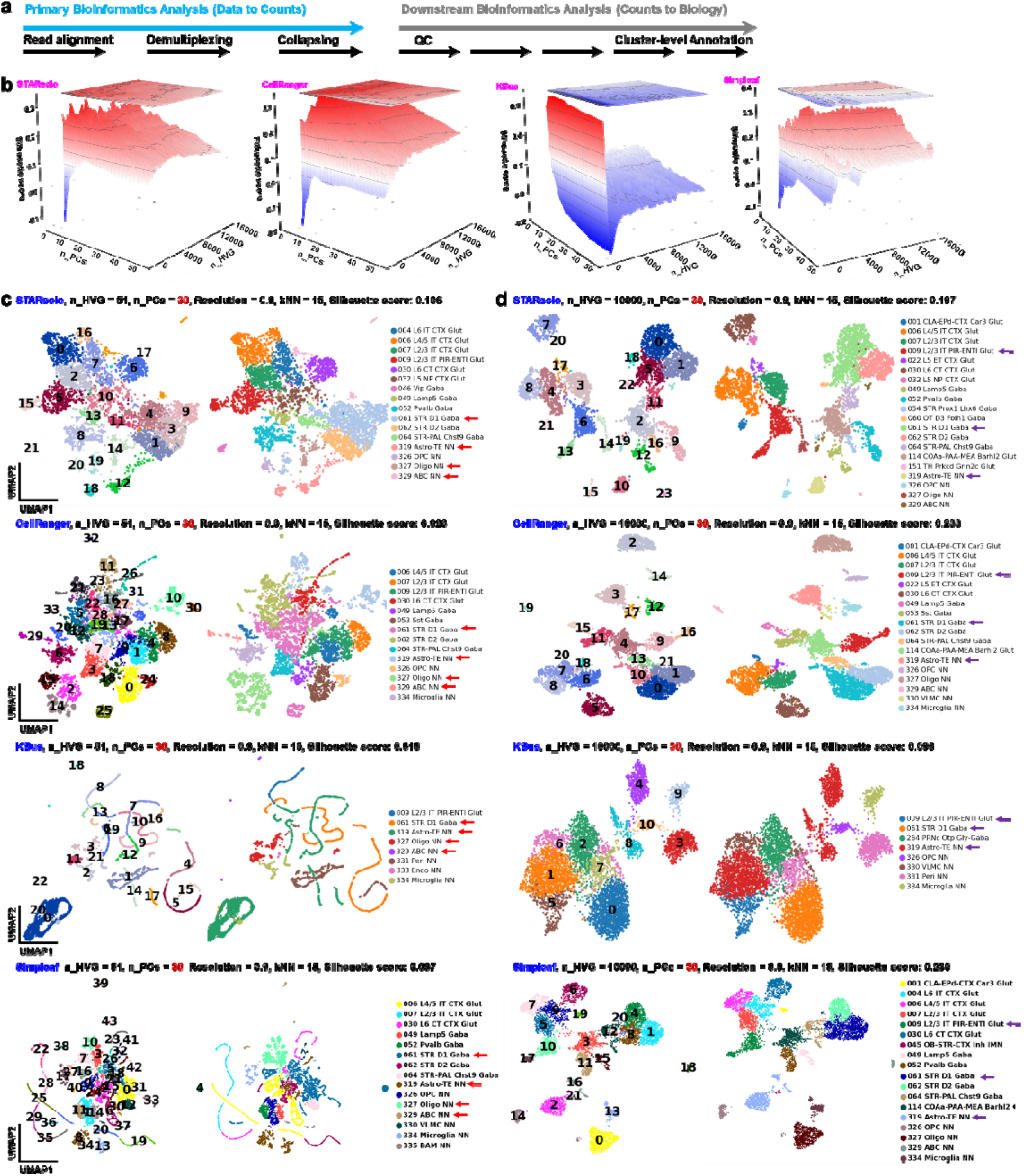
Impact of primary processing and downstream parameters on cluster-level cell type annotation. **a**) Schematic of the end-to-end scRNA-seq evaluation workflow. The process begins with raw read processing via one of four primary mapping pipelines (STARsolo, Cell Ranger, KBus, or Simpleaf) to generate a gene-by-cell count matrix. This matrix is then processed through a standardized downstream workflow containing key tunable parameters, notably the number of highly variable genes (n_HVG) and principal components (n_PCs). Final biological interpretation relies on cluster-level annotation, determined by first grouping cells via the Leiden algorithm and subsequently assigning a consensus cell type to each cluster based on a majority vote of individual cell predictions. **b**) 3D surface plots visualizing parameter optimization for clustering. The z-axis displays the Silhouette Score-a metric of cluster cohesion and separation-resulting from a simultaneous grid search over varying n_PCs (x-axis) and n_HVGs (y-axis). Conducted on the 10x Genomics 5k Adult Mouse Brain dataset for each primary processing method, this analysis identifies parameter plateaus that yield the most robustly separated clusters. **c-d**) Comparison of cluster-level annotation outcomes across distinct downstream parameter regimes. For all pipelines, clustering was performed using a fixed Leiden resolution of 0.9 and k=15 nearest neighbors. **c**) Under a highly restrictive feature selection setting (n_HVG = 51, n_PCs = 30), five common cell types were consistently identified across all four primary pipelines (highlighted by arrows), with respective Silhouette Scores displayed. (d) Conversely, utilizing a much broader feature space (n_HVG = 10,000, n_PCs = 30) resulted in only three cell types being universally identified across all methods. This contrast illustrates the acute sensitivity of final biological annotations to the interplay between primary processing and downstream parameter choices.

When n_HVG was increased to 10,000 while keeping n_PCs = 30, the resulting Leiden clustering and cell annotations changed markedly (Fig.□2d). Under these broader, more optimal conditions, STARsolo, Cell Ranger, and Simpleaf all capitalized on the expanded feature space to resolve a high-resolution landscape of discrete cell identities. Each of these three pipelines identified an expanded repertoire of 18 specific cell types. Furthermore, their clusters demonstrated exceptional internal consistency; for instance, STARsolo achieved a CAS of 82.75% for STR D1 Gaba, while Simpleaf reached 90.60% for L4/5 IT CTX Glut and 100% for CLA-EPd-CTX Car3 Glut. (Simpleaf’s Silhouette Score under these optimal conditions correspondingly improved to 0.230). Conversely, KBus completely failed to utilize the expanded feature set. It stubbornly collapsed the data into only 8 highly mixed clusters with persistently abysmal purity, never exceeding a 42% match rate and frequently dropping below 20% (e.g., 9.84% CAS for PRNc Otp Gly-Gaba). Overall, these results demonstrate that both the primary processing pipeline and downstream parameter choices independently shape the final clustering and cell type annotation. Crucially, they reveal that while lightweight mapping can be highly effective-as evidenced by Simpleaf’s strong performance matching alignment-based methods-poor algorithmic implementation at the primary processing stage (as seen with KBus) can inflict irreversible damage on the data structure that no amount of downstream parameter tuning can rescue.

### Impact of primary data processing on direct cell-level annotation and downstream visualization

In conventional scRNA-seq analysis, cell identities are often inferred through a cluster-centric workflow in which final cell-type assignments may be sensitive to downstream analytical parameters, including the number of highly variable genes (HVGs) and principal components (PCs). To distinguish the effects of primary data processing from those introduced during downstream analysis, we developed a direct cell-level annotation strategy (Fig. 3a). Raw sequencing reads from the 10x Genomics 5k Adult Mouse Brain dataset were processed using four independent pipelines-STARsolo, Cell Ranger, KBus, and Simpleaf-to generate distinct gene-by-cell count matrices. Following standard quality control, each cell was annotated individually using its complete transcriptomic profile before dimensionality reduction. This approach established a set of cell-identity labels for each primary-processing pipeline and enabled the effects of different analytical stages to be evaluated separately.

**Fig. 3.**
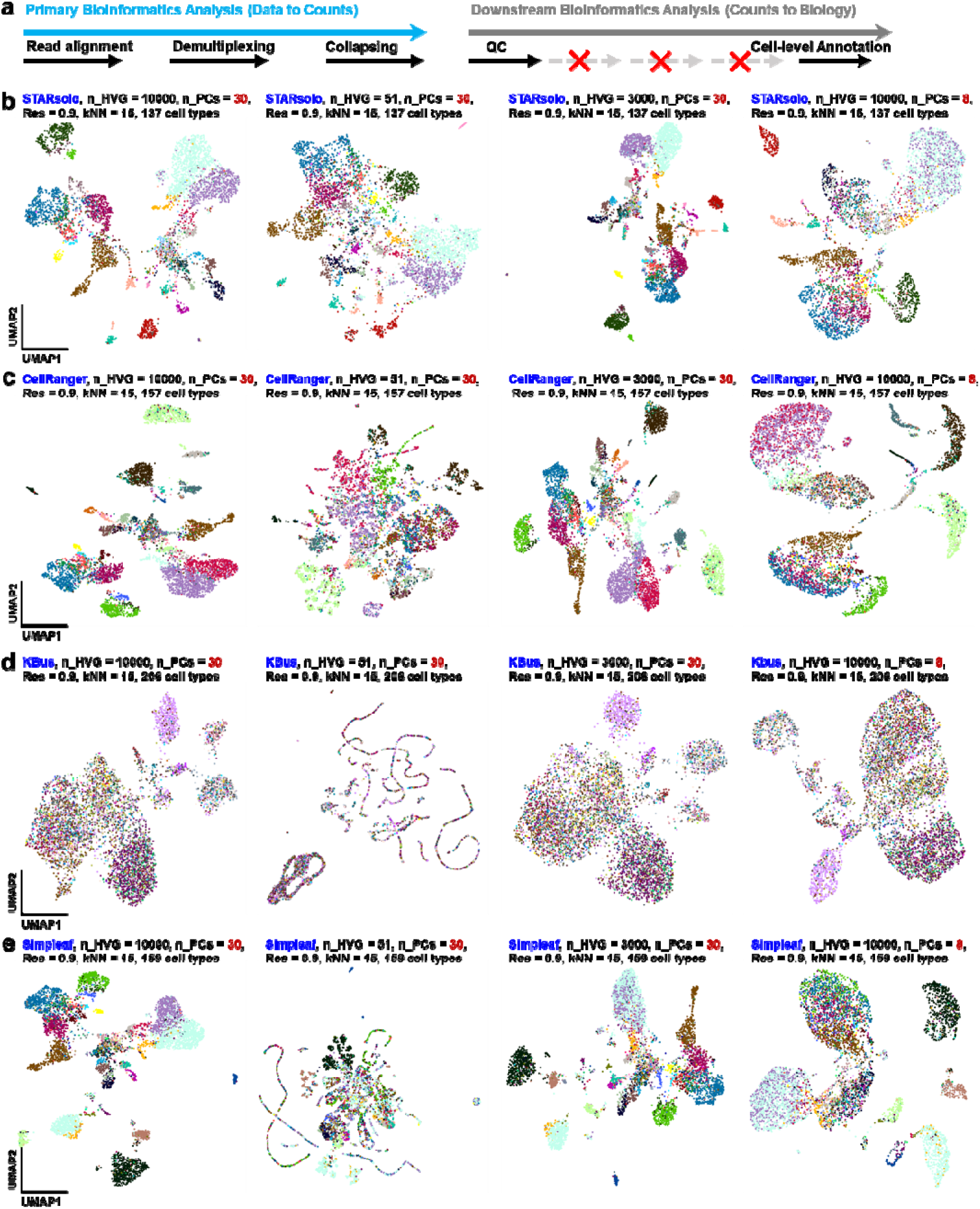
Influence of primary data processing on direct cell-level annotation and spatial embedding in the 10x Genomics 5k Adult Mouse Brain dataset. **a**) Schematic of the direct cell-level annotation workflow. In contrast to cluster-consensus approaches, this pipeline generates a primary gene-by-cell matrix via one of four methods (STARsolo, Cell Ranger, KBus, or Simpleaf), applies standard quality control filtering, and independently annotates each cell based on its distinct gene expression profile. This biological identity assignment is calculated once per primary method and remains fixed across all subsequent dimensionality reduction steps. **b–e**) Impact of downstream parameter variations on UMAP embeddings of pre-annotated cells. For each primary processing pipeline-(b) STARsolo, (c) Cell Ranger, (d) KBus, and (e) Simpleaf-the fixed, cell-level annotations (indicated by color) are projected onto UMAP spaces generated under four distinct downstream parameter regimes (n_HVG/n_PCs pairs: 51/30, 3000/30, 10000/30, and 10000/8). These projections visually demonstrate how the selection of highly variable genes and principal components fundamentally restructures the spatial topology and perceived grouping of the dataset, despite the underlying biological identities of the cells remaining strictly constant.

We then assessed whether downstream parameter variation altered the direct cell-level annotations or only the visualization of the annotated populations. For each primary-processing pipeline, UMAP embeddings were generated using four parameter combinations: 51 HVGs with 30 PCs, 3,000 HVGs with 30 PCs, 10,000 HVGs with 30 PCs, and 10,000 HVGs with 8 PCs (Fig. 3b-e and Supplementary Figs. S4-S7). The clustering resolution and k-nearest-neighbor parameters were held constant at 0.9 and 15, respectively. Although the spatial organization and apparent separation of cells varied across these settings, the direct cell-level annotations remained consistent within each pipeline. STARsolo yielded 137 cell types in all four parameter settings, Cell Ranger yielded 157, KBus yielded 206, and Simpleaf yielded 159. Thus, changing the number of HVGs from 51 to 3,000 or 10,000, or reducing the number of PCs from 30 to 8, did not alter the cell-type assignments established before dimensionality reduction. These results indicate that downstream parameters can substantially affect embedding geometry while leaving the underlying direct annotations unchanged.

The quality of the primary count matrices nevertheless influenced how clearly the annotated populations were represented in the embeddings. For STARsolo, Cell Ranger, and Simpleaf, the directly annotated cell populations generally formed relatively cohesive and distinguishable groups under the more favorable parameter settings, suggesting that these matrices retained transcriptomic structure sufficient to resolve multiple cellular populations. In contrast, the KBus-derived matrix showed greater mixing among annotated populations and less distinct spatial organization in the UMAP representations. This observation also highlights that the number of directly annotated cell types is not, by itself, a sufficient measure of data quality or biological resolution. In particular, KBus produced the largest number of annotations, but these annotations were not necessarily accompanied by clear separation in the low-dimensional embeddings.

Primary-processing methods also differed in the biological and technical content retained from the sequencing reads. Although a core set of cell types was shared across pipelines, each method produced a distinct cell-type recovery profile. Importantly, these profiles were stable across all four downstream parameter combinations: STARsolo consistently yielded 137 cell types, Cell Ranger 157, KBus 206, and Simpleaf 159. The number of annotated cell types therefore reflected differences established during primary mapping and quantification rather than changes in the selected HVG or PC parameters. Simpleaf produced a cell-type repertoire similar in size to that of Cell Ranger in this dataset, while STARsolo yielded fewer and KBus yielded more directly annotated cell types. These differences should be interpreted together with the cohesion and separation of the corresponding cell populations in the embeddings rather than as standalone measures of pipeline performance.

Similarly, the technical overlap among pipelines-representing the specific cell barcodes retained after quality-control filtering-varied according to the primary-processing method. Because barcode retention and direct cell-level annotation were determined before dimensionality reduction, these pipeline-specific features were not expected to change with the downstream HVG or PC settings. Consequently, the results distinguish two separable sources of variability in the scRNA-seq workflow: (i) differences in cell capture, barcode retention, and gene quantification introduced during primary data processing; and (ii) changes in the geometry and visual organization of cell populations caused by downstream dimensionality-reduction parameters.

By applying cell-level annotation before dimensionality reduction, this framework separates changes in biological assignment from changes in visualization. The consistent annotations observed across all tested parameter settings demonstrate that direct cell-level labels provide a stable reference for comparing primary-processing methods, while the differing UMAP structures reveal how the quality and properties of the underlying count matrices influence downstream representation. Overall, these findings emphasize that primary data processing is a major determinant of the recovered biological repertoire and its apparent organization, whereas downstream parameter selection primarily affects the geometry through which those populations are visualized.

### A consistency metric reveals pipeline- and parameter-dependent differences in cell-type annotation concordance

The choice of primary-processing pipeline shapes the count matrix used for downstream clustering and cell-type annotation. To assess how these differences relate to final annotations, we established a CAS framework (Fig. 4a). The framework compares direct cell-level annotations, in which cells are classified individually from their transcriptomic profiles, with cluster-level consensus annotations, in which each Leiden cluster receives the majority label of its constituent cells. CAS is the fraction of cells in a cluster whose direct label matches that consensus label. Its distribution across clusters describes cell-to-cluster annotation concordance, rather than the biological accuracy of the assigned labels.

**Fig. 4.**
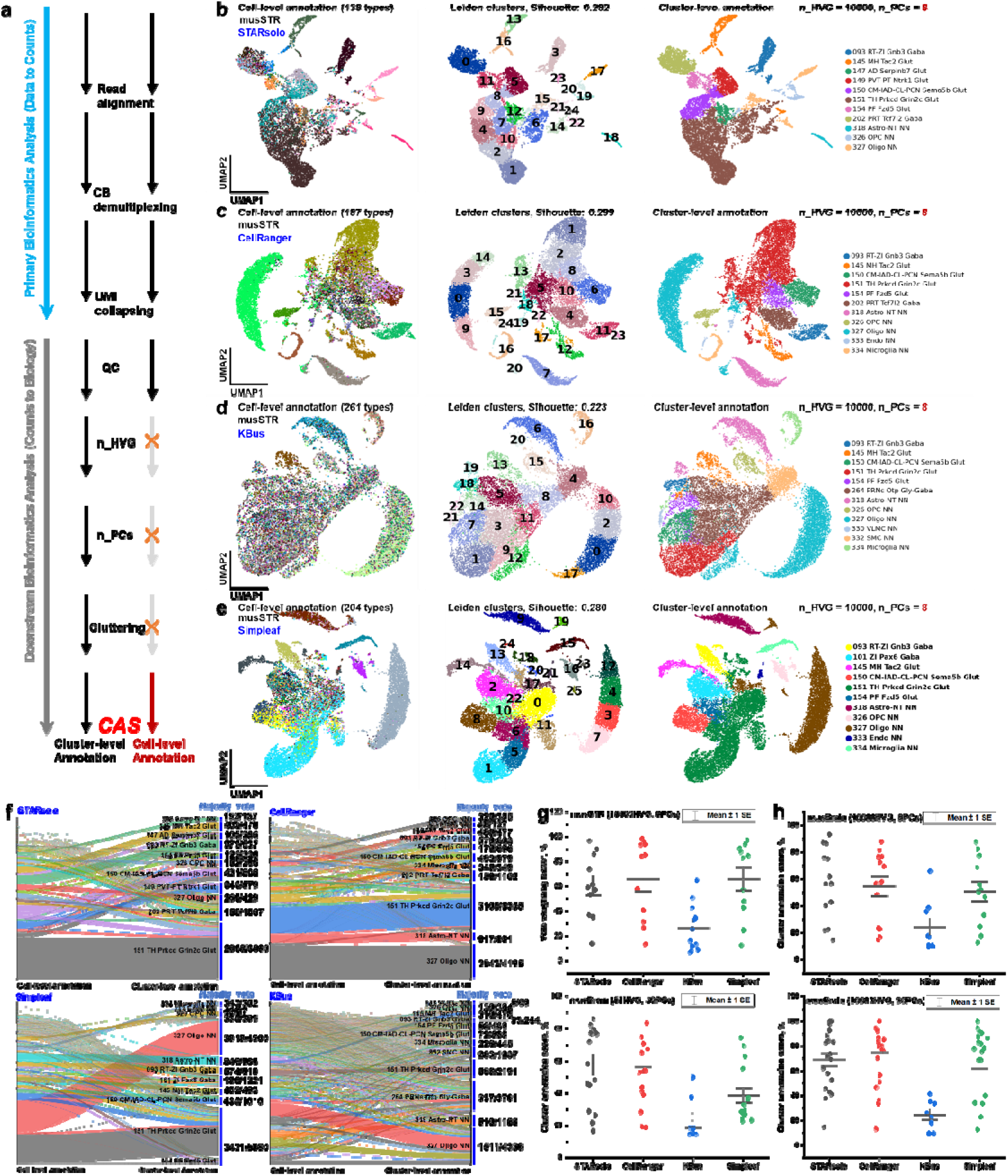
Benchmarking annotation consistency across pipelines with the CAS. **a**) A schematic of the annotation consistency assessment framework. The method leverages two outputs: direct cell-level annotations (where each cell is labeled independently) and cluster-level consensus annotations (where a Leiden cluster is assigned a label by a majority vote of its constituent cells). To quantify consistency, we define a CAS for each Leiden cluster as the fraction of cells within that cluster whose individual cell-level label matches the final cluster-level consensus label. A higher score indicates stronger agreement between the granular and grouped annotations. An overall performance metric for each pipeline is derived from the distribution of these scores across all clusters. **b-e**) Visualization of the annotation framework components on an in-house single-nucleus RNA-seq dataset from wild type mouse striatum (musSTR), processed with n_HVG=10000 and n_PCs=8. For each primary method-(**b**) STARsolo, (**c**) CellRanger, (**d**) KBus, and (**e**) Simpleaf-three UMAPs are shown: the direct cell-level annotations, the Leiden cluster partitions, and the resulting cluster-level consensus annotations. **f**) Quantitative visualization of annotation concordance. Alluvial plots show the flow of cells from their initial cell-level annotation (left) to their final cluster-level consensus label (right). The width of each stream is proportional to the number of cells, providing a direct visual measure of how consistently cell populations are classified at both scales. The total number of cells assigned to each consensus label is noted. **g**) Comparison of annotation consistency across pre-processing pipelines on the musSTR dataset. The scatter-and-interval plot displays the distribution of Cluster Annotation Scores (CAS) for clusters generated by four different methods: STARsolo, CellRanger, KBus, and Simpleaf. Each point represents an individual cluster. The CAS reflects the percentage of cells within a cluster that match the consensus label, serving as a metric for annotation purity and internal consistency. This allows for a direct evaluation of the cell-to-cluster annotation concordance produced by each pipeline. **h**) Generalization and robustness of benchmarking results. To validate the consistency analysis, the evaluation was extended to the independent 10x Genomics 5k Adult Mouse Brain dataset. These three plots compare the distribution of Cluster Annotation Scores (CAS) for STARsolo, CellRanger, KBus, and Simpleaf under three distinct downstream clustering parameter settings (varying combinations of highly variable genes [n_HVG] and principal components [n_PCs]). This comparison demonstrates how annotation concordance varies across pipelines and downstream parameter settings.

We first applied this framework to an in-house single-nucleus dataset of wild-type mouse striatum (musSTR), using 10,000 highly variable genes (HVGs) and 8 principal components (PCs) (Supplementary Fig. S8). The paired views of direct annotations, Leiden partitions, and consensus annotations show how each pipeline organizes annotated populations (Fig. 4b-e). STARsolo and Cell Ranger generally produced distinct populations that corresponded to coherent clusters, whereas KBus showed greater mixing. Simpleaf did not follow the same pattern as KBus: several of its annotated populations also mapped coherently to clusters, although the correspondence varied by cell type. The alluvial plots complement these views by showing how cells move between direct and consensus labels (Fig. 4f); the CAS distributions quantify the agreement within clusters (Fig. 4g).

The musSTR CAS values make the distinction between the two lightweight-mapping workflows particularly clear. Simpleaf achieved scores of 91.14% for Oligo, 87.89% for Astro-NT, 94.10% for RT-ZI Gnb3 Gaba, and 97.44% for Microglia, close to the corresponding Cell Ranger scores of 93.99%, 93.48%, 95.85%, and 98.85%. KBus scored substantially lower for these shared populations. Simpleaf was nevertheless uneven: its scores fell to 12.78% for ZI Pax6 Gaba and 24.93% for PF Fzd5 Glut. Thus, Simpleaf approached the alignment-based pipelines for several populations, but the data do not support treating its performance as uniformly equivalent to theirs. KBus had low scores for many listed populations, including 8.96% for PRNc Otp Gly-Gaba and 8.11% for CM-IAD-CL-PCN Sema5b Glut.

The pipelines also differed in the number of reported cell types: 138 for STARsolo, 187 for Cell Ranger, 261 for KBus, and 227 for Simpleaf (Supplementary Figs. S9-S12). These counts cannot independently distinguish improved biological resolution from over-separation. In particular, KBus combined a larger reported repertoire with low CAS for many clusters, whereas Simpleaf’s larger repertoire co-occurred with high concordance for several populations. The additional reported types therefore require evaluation beyond their number alone.

We next examined a corresponding Huntington’s disease mouse-model dataset under 8-PC and 30-PC settings (Supplementary Figs. S13-S14). Pipeline-specific differences in annotation concordance remained apparent, and changing the number of PCs altered the resulting cluster-level assignments. These comparisons show that downstream parameter choices can affect the relationship between direct cell labels and cluster consensus labels, in addition to any differences introduced during primary processing.

Finally, we evaluated an independent 10x Genomics 5k Adult Mouse Brain dataset under three HVG/PC settings (Fig. 4h). At 10,000 HVGs and 30 PCs, Simpleaf achieved CAS values of 87.62% for Oligo, 86.59% for L2/3 IT PIR-ENTl Glut, and 83.01% for STR D2 Gaba, close to the corresponding Cell Ranger values of 85.22%, 84.39%, and 80.28%. KBus scored 33.82% for L2/3 IT PIR-ENTl Glut in the same setting. Yet Simpleaf’s score was only 12.95% for COAa-PAA-MEA Barhl2 Glut. Its variation was also evident in the other settings: at 10,000 HVGs and 8 PCs, scores ranged from 87.73% for Oligo to lower values for several neuronal populations; at 51 HVGs and 30 PCs, Oligo scored 74.47% while L4/5 IT CTX Glut scored 22.70%.

Together, these results indicate that STARsolo and Cell Ranger generally maintained strong cell-to-cluster annotation concordance, while KBus showed low concordance for many of the reported populations. Simpleaf often performed substantially better than KBus and matched or exceeded the alignment-based pipelines for selected populations, but its results were more dependent on cell type and analytical setting. CAS therefore helps locate discrepancies between direct and consensus annotations; establishing which pipeline yields the most biologically accurate cell types requires independent evidence.

## Discussion

We evaluated how four primary-processing workflows-STARsolo, Cell Ranger, KBus, and Simpleaf-relate to downstream interpretation of scRNA-seq and snRNA-seq data. Using simulated and empirical datasets, we examined technical performance, clustering outcomes, and agreement between direct cell-level and cluster-level consensus annotations. We also introduced the Marker Concordance Score (MCS) to assess marker-gene prevalence within annotated populations. These measures address different aspects of a pipeline’s output: technical performance does not necessarily predict downstream annotation concordance, and neither CAS nor MCS independently establishes the biological correctness of a cell-type label.

Across the empirical analyses reported here, STARsolo and Cell Ranger generally showed strong cell-to-cluster annotation concordance. KBus had low CAS for many of the reported populations, including under settings in which the other pipelines achieved higher scores. Simpleaf, however, should not be grouped with KBus as a uniformly low-concordance lightweight workflow. It approached or exceeded the alignment-based pipelines for selected populations in both the musSTR and 5k Adult Mouse Brain datasets. Its scores were also low for other populations, particularly under some HVG/PC settings. The principal distinction is therefore not a simple ranking of alignment-based over lightweight methods, but variation among workflows and across cell types and analytical conditions. The larger number of cell types reported by KBus-and, to a lesser extent, Simpleaf-cannot by itself be interpreted as either improved resolution or over-separation.

Primary-processing differences were reflected in downstream results, but their effects must be interpreted alongside downstream analysis choices. Changing n_HVG or n_PCs altered embedding geometry, clustering, and the resulting cluster-level consensus annotations. Direct cell-level labels, assigned before dimensionality reduction, remained fixed within each pipeline across those parameter settings. This design distinguishes changes in cluster-level representation from changes in the direct labels; it does not, by itself, isolate which primary-processing operation caused a difference between pipelines. Differences in barcode retention, read assignment, and quantification may all contribute, and their individual effects require more targeted tests.

The parameter analyses further show why no single clustering metric provides a complete assessment. Silhouette Score describes the separation and cohesion of clusters in a chosen representation, whereas CAS measures agreement between direct labels and cluster consensus labels. A well-separated cluster need not correspond to a biologically accurate cell type, and a high CAS can occur when the direct and consensus annotations agree on an incorrect label. Parameter selection should therefore be assessed using complementary evidence rather than a Silhouette Score or CAS threshold alone. The low-feature setting was particularly informative in this respect: it produced substantially different outcomes across pipelines, while Simpleaf retained relatively high concordance for some populations but not others.

Several limitations qualify these findings. CAS depends on the direct annotation method, the cell types present, and the clusters produced under each setting; comparisons are also affected when pipelines retain different sets of cells. Likewise, marker-gene prevalence can support an assessment of within-population coherence but is not an independent ground truth for cell identity. The results presented here do not establish that the pipeline differences arise solely from mapping strategy, nor do they support a universal ranking of biological accuracy across datasets. In particular, the observed variation in Simpleaf CAS argues against describing STARsolo, Cell Ranger, and Simpleaf as uniformly equivalent or uniformly robust to parameter changes. Computational efficiency is an important consideration, but claims about relative speed or resource use should be supported by the corresponding benchmark results.

## Methods

### Collection of tissue samples

B6-hHTTCAG130 mice (Strain No. T054804, GemPharmatech), which harbor a human HTT gene fragment containing 130 CAG repeat expansions, were used as a Huntington’s disease model. Striatal tissues from these mice were collected and designated as musHuntingtonD-STR, while corresponding striatal samples from wild-type control mice were designated as musWT-STR. All animals were maintained under specific pathogen-free (SPF) conditions, provided with autoclaved food, and given acidified, autoclaved water. All experimental procedures involving animals were performed in strict accordance with institutional guidelines and approved by the Animal Care and Use Committee of Shenzhen University of Advanced Technology.

### snRNA-seq Library preparation

Single nuclei were isolated from the collected mouse striatum samples, and snRNA-seq libraries were prepared using the 10x Genomics Chromium Next GEM Single Cell 3’ v3.1 Kit according to the manufacturer’s protocol. In this workflow, individual nuclei were encapsulated in gel bead-in-emulsions (GEMs), lysed to release mRNA, reverse-transcribed into cDNA, and subsequently pre-amplified for downstream sequencing.

### Sequencing

The purified PCR libraries were submitted for sequencing using either the Illumina NovaSeq 6000 (PE150) platform or the MGISEQ-2000 (PE150) platform. The data output was generated according to the specified yield set.

### Read processing and gene assignment

Etiam sit amet euismod ipsum, eu pretium neque. Maecenas et maximus tortor. Ut laoreet ullamcorper tincidunt. Aenean dapibus ullamcorper sapien, eu iaculis nisl aliquet et. Vivamus ornare, dui ultricies cursus euismod, odio magna porttitor diam, quis dictum metus ante vel metus. Class aptent taciti sociosqu ad litora torquent per conubia nostra, per inceptos himenaeos.

The initial stage of scRNA-seq processing assigns each raw sequencing read to its gene of origin, a step that defines the fundamental trade-off between base-level mapping precision and computational speed. The four benchmarked pipelines fall into two distinct strategic groups: those based on spliced genomic alignment and those based on k-mer mapping.

#### 1. Spliced Genomic Alignment (Cell Ranger, STARsolo)

Both Cell Ranger and STARsolo utilize the same core algorithm, STAR, to perform spliced alignment of reads against a reference genome. This method identifies the precise genomic locus (chromosome, start, end, and splicing pattern) for each read. A read is subsequently assigned to a gene if its aligned coordinates overlap with the annotated exonic boundaries of that gene.

The gene assignment rule can be formalized as follows: Let a read *r* be mapped to a genomic locus *L*. Let the set of exons for gene g be E_*g*_. The assignment function is:

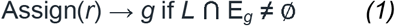

Where:

- Assign(r)→g indicates that the function Assign takes read rr as input and returns gene *g* as the gene to which *r* is assigned.
- L is the set of genomic coordinates covered by the aligned read, including split intervals in the case of splicing.
- E_*g*_ is the set of exon intervals annotated for gene *g*.
- ∩ denotes the set intersection operation-in this context, it tests whether the aligned portion(s) of *r* share any overlapping genomic coordinates with an annotated exon.
- ≠ ∅ means “is not an empty set”; here, it requires that the intersection contains at least one nucleotide of overlap between LL and E_*g*_.
- ∅ is the empty set symbol, representing “no overlap”. Symbolically: Read → [Genomic Alignment] → Gene

Where the arrow (→) denotes the transformation of raw read sequences into gene-level assignments through the alignment process.

#### 2. K-mer-based Mapping

This approach bypasses computationally intensive base-to-base alignment by decomposing reads into constituent k-mers and matching them against a pre-built index of the transcriptome. However, the specific algorithms for this matching differ.

##### 2.1 KBus (Pseudoalignment)

KBus uses a pseudoalignment approach to assign sequencing reads to transcripts without performing full base-to-base genomic alignments. Instead, each read *r* is decomposed into a set of fixed-length subsequences called *k*-mers. These *k*-mers are matched to paths in a colored de□Bruijn graph, which is constructed from the reference transcriptome and encodes all known transcript sequences and their shared subsequences. The output of this search is the set of transcripts whose *k*-mers are fully compatible with the *k*-mers of *r*. This set is known as the Transcript Compatibility Class (TCC) or, equivalently, the equivalence class (*EC*) of the read. The *EC* represents all transcripts from which *r* could plausibly originate.

To generate gene-level counts, each equivalence class is resolved to one or more genes based on the gene annotations associated with the transcripts in that *EC*. The rule for gene assignment can be expressed formally as:

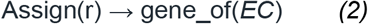

Where:

- Assign(*r*)→gene_of(*EC*) indicates that the function *Assign* takes the read *r* as input, determines its equivalence class *EC* via pseudoalignment, and outputs gene_of(*EC*), i.e., the gene or genes corresponding to the transcripts in *EC*.
- *r* is the sequencing read being assigned.
- *EC* is the equivalence class (Transcript Compatibility Class) containing all transcripts whose sequence composition matches the *k*-mers of *r*.
- gene_of(*EC*) is the function that maps an equivalence class to the corresponding gene(s) in the reference annotation.
- Pseudoalignment refers to the process of determining *EC* without computing exact genomic coordinates, using *k*-mer matching in the de□Bruijn graph.
- “Equivalence Class” denotes the transcript set definition-all members of *EC* are indistinguishable for *r* given the available sequence information.

Symbolically: Read → [Pseudoalignment | Equivalence Class] → Gene

Here, the vertical bar (∣) separates the two conceptual steps: pseudoalignment generates the equivalence class *EC*, which is then collapsed to its corresponding gene(s). The arrow (→) denotes the transformation from a raw read sequence to a gene assignment through this two-stage process.

##### 2.2 Alevin-fry (Quasi-mapping)

Alevin-fry applies a quasi-mapping strategy to associate sequencing reads with transcripts and their approximate positions without performing full alignment. For each read *r*, the algorithm decomposes it into *k*-mers and uses a highly optimized index-combining a suffix array and a hash table-to rapidly identify the minimal set of transcripts and positions from which the read’s sequence could plausibly derive. This location-aware method differs from pseudoalignment by explicitly storing and considering transcript positional information, allowing it to determine not only which transcripts are compatible but also the likely genomic or transcript coordinate range for each match.

From this process, Salmon produces a set of quasi-mapping loci, *M*, where each element is a pair (*t, pos*) consisting of transcript t and position *pos* within that transcript. The set of compatible transcripts is then defined as:

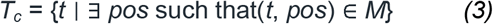

The final gene assignment rule is:

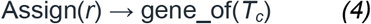

Where:

- Assign(*r*)→gene_of(*T*_*c*_) indicates that the function *Assign* takes read *r* as input, determines the set of compatible transcripts *T*_*c*_ via quasi-mapping, and outputs gene_of(*T*_*c*_), i.e., the gene or genes associated with those transcripts.
- *r* is the sequencing read under consideration.
- *M* is the set of quasi-mapping loci for *r*, each represented as a (*t, pos*) pair, where *t* is a transcript identifier and *pos* is the alignment position within *t*.
- *Tc* (“transcript compatibility set”) is the set of all transcripts *t* for which there exists at least one *pos* with (*t, pos*) ∈ *M*. The existential quantifier ∃ (“there exists”) specifies that a transcript is included if at least one matching position is found.
- gene_of(*T*_*c*_) is a function mapping the transcript set *T*_*c*_ to its corresponding gene(s) based on the reference annotation.
- Quasi-mapping refers to finding transcript and position pairs with high probability based on *k*-mer matches without performing a base-by-base alignment.
- The vertical bar (∣) in the symbolic representation separates the two major conceptual steps: quasi-mapping to determine locus IDs, and collapsing those loci to gene assignments.
- The arrow (→) denotes transformation from the raw read to a gene assignment through this process.

Symbolically: Read → [Quasi-mapping | Locus ID] → Gene

Here, *Quasi-mapping* derives the mapping loci (*t, pos*) and *Locus ID* resolution determines *T*_*c*_, which is then collapsed to its gene-level assignment.

### Cell barcode demultiplexing

Following read-to-gene assignment, the next critical stage is cell barcode demultiplexing, which resolves raw barcode sequences into a final set representing individual cells. This process has two coupled objectives: first, identifying the subset of barcodes corresponding to genuine cell-containing droplets (cell calling), and second, correcting sequencing or amplification errors in observed barcodes by assigning them to a true barcode from the called set. The four benchmarked pipelines approach these objectives with distinct algorithms and philosophies.

#### 1. Cell Ranger (Integrated Probabilistic Model)

Cell Ranger uses an integrated probabilistic framework to perform cell calling and barcode correction in a single step. The process starts from a large set of barcodes observed in sequencing data, many of which originate from background (ambient) RNA rather than actual cells. To identify genuine cell-associated barcodes, Cell Ranger analyzes the UMI count distribution across all observed barcodes. Barcodes whose UMI counts significantly exceed the expected background model are designated as “high-confidence” cell barcodes, forming the set *C*, also known as the whitelist.

In real sequencing data, sequencing errors may produce observed barcodes (*b*_obs_) containing mismatches to their true underlying barcodes (*b*_true_). Cell Ranger corrects these errors within a Bayesian statistical model. The model evaluates all candidate barcodes in *C* as possible sources of *b*_obs_, using the per-base sequencing quality scores *Q* to determine the probability that the observed mismatches arose from sequencing errors.

The correction rule selects the maximum a posteriori (MAP) estimate:

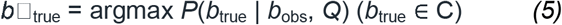

Where:

- *b*□_true_ denotes the final corrected barcode, representing the highest⍰probability true barcode for the given observed barcode.
- *b*_true_ refers to a candidate “true” barcode from the whitelist *C*.
- *b*_obs_ represents the barcode sequence observed in the sequencing data, which may contain errors introduced during sequencing.
- *Q* indicates the per⍰Jbase sequencing quality scores, reflecting the probability that each base call is correct.
- *C* designates the whitelist containing all high⍰confidence cell barcodes identified during cell calling.
- argmax specifies the mathematical operator that returns the *b*_true_ value which maximizes the posterior probability.
- *P*(*b*_true_∣*b*_obs_, *Q*) expresses the posterior probability that the observed barcode *b*_obs_ was derived from the candidate true barcode *b*_true_, given the estimated sequencing error profile from *Q*.

Symbolically: b? → [P(Q) | Whitelist] → b⍰

Where:

- *b*_?_ denotes the observed barcode of uncertain accuracy.
- *P*(*Q*) indicates the probability model derived from base quality scores, estimating the likelihood of sequencing errors at each base position.
- Whitelist refers to the set *C* of high⍰Jconfidence barcodes determined during cell calling.
- *b*⍰represents the final corrected barcode selected using the Maximum A Posteriori (MAP) estimation.

#### 2. STARsolo (Deterministic Whitelist Correction)

STARsolo determines cell barcodes in two stages: cell calling and error correction, with each stage handled separately.

In the cell calling phase, STARsolo first identifies a preliminary set of barcodes associated with cells. This can be done by:

- Ranking all observed barcodes by their total UMI counts, then identifying an inflection point (“knee-point”) separating high-count (cell) barcodes from low-count (background) barcodes.
- Alternatively, applying the EmptyDrops algorithm to statistically distinguish real cells from droplets containing ambient RNA.

For error correction, STARsolo takes a deterministic approach. It uses the manufacturer-provided whitelist *W*, which contains all valid barcode sequences predefined for the specific chemistry kit (e.g., 10x Genomics v3 chemistry). The observed barcode *b*_obs_ is compared to each whitelist barcode *b*_*w*_ using the Hamming distance *H*, which measures the number of nucleotide positions at which the two sequences differ.

The correction rule is based on a fixed Hamming distance threshold to the predefined whitelist:

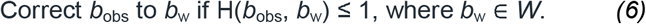

Where:

- *b*_obs_ denotes the observed barcode sequence from the sequencing run, which may contain sequencing errors.
- *b*_*w*_ refers to a barcode sequence in the manufacturer’s whitelist *W*.
- *W* represents the complete, predefined whitelist of valid barcode sequences for the library preparation chemistry used.
- *H*(*b*_obs_,*b*_*w*_) indicates the Hamming distance between *b*_obs_ and *b*_*w*_, i.e., the number of single-nucleotide mismatches.
- ≤1 specifies the threshold allowing barcode correction if the observed barcode differs by at most one base from exactly one whitelist barcode.
- Correction condition describes the rule that if no whitelist barcode matches within a Hamming distance of 1, or if more than one whitelist barcode matches at that distance, *b*_obs_ remains uncorrected to prevent ambiguity.

Symbolically: b? → [H≤1 | Whitelist] → b⍰

Where:

- *b*_?_ denotes the observed barcode to be validated.
- *H* ≤ 1 specifies the Hamming distance threshold of one or fewer mismatches.
- Whitelist refers to the valid barcode set *W*.
- *b*⍰ represents the corrected barcode, which exactly matches one whitelist entry within the given threshold.

#### 3. KBus (Uniqueness-Constrained Whitelist Correction)

KBus performs barcode correction using a deterministic approach similar to STARsolo, but with an additional uniqueness constraint to increase assignment specificity. In the first step, KBus uses the predefined manufacturer’s whitelist *W*, which contains all valid barcode sequences for the library preparation chemistry. Each observed barcode bobs*b*obs from the sequencing run is compared to every whitelist barcode *b*_*w*_ by calculating the Hamming distance *H*(*b*_obs_, *b*_*w*_), defined as the number of single-nucleotide positions where the sequences differ.

A candidate correction is considered only if the Hamming distance from bobs*b*obs to at least one *b*_*w*_ in *W* is ≤□1. However, KBus enforces a strict uniqueness requirement: there must exist **e**xactly one whitelist barcode within this distance. This is expressed by the logical quantifier ∃! (pronounced “there exists exactly one”). If two or more whitelist barcodes meet the Hamming distance condition, or if no match exists, *b*_obs_ remains uncorrected to avoid ambiguity and misassignment.

The barcode correction rule is:

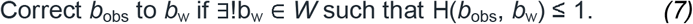

Where:

- *b*_obs_ denotes the observed barcode sequence from the sequencing output, potentially carrying sequencing errors.
- *b*_*w*_ refers to a valid barcode sequence found in the manufacturer’s whitelist *W*.
- *W* represents the complete, predefined whitelist of valid barcode sequences for the specific library preparation chemistry.
- *H*(*b*_obs_, *b*_*w*_) indicates the Hamming distance between *b*_obs_ and *b*_*w*_, equal to the number of base positions that differ.
- ≤1 specifies the maximum distance allowed for considering a correction, i.e., one or fewer mismatches.
- ∃! denotes the logical statement “there exists exactly one,” which is the uniqueness condition that must be satisfied for correction.

Symbolically: b? → [H≤1, Unique!] → b⍰

Where:

- *b*_?_ denotes the observed barcode to be validated.
- *H*≤1 specifies the maximum allowable Hamming distance for correction.
- Unique! indicates that exactly one whitelist barcode satisfies the Hamming distance condition.
- *b*⍰ represents the final corrected barcode, guaranteed to match exactly one whitelist entry within the specified threshold.

#### 4. Alevin-fry (Data-Driven Permit List Correction)

Unlike Cell Ranger, STARsolo, or KBus, which rely on a predefined manufacturer’s whitelist, Alevin-fry uses a data-driven, adaptive approach to determine valid cell barcodes. The process begins by scanning all observed barcodes *b* from the sequencing data and calculating their total UMI counts. These counts are plotted as a histogram ordered from the highest count to the lowest.

Alevin-fry then applies an algorithm-such as knee-point detection or mixture modeling**-**to determine a UMI count threshold θ that best separates real cell barcodes from background (ambient) RNA. Any barcode with a count greater than θθ is included in the permit list *P*, which acts as the set of trusted barcodes for that particular dataset. Unlike a fixed whitelist *W, P* is empirically derived from the data.

The permit list is defined as:

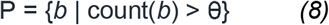

Once *P* is established, the observed barcode *b*_obs_ is compared against the entries *b*_p_ ∈ *P* using a distance criterion (often Hamming distance ≤□1). If exactly one barcode in *P* meets the distance requirement, *b*_obs_ is corrected to that *b*_p_. If no match or multiple matches exist within the threshold, *b*_obs_ remains uncorrected to avoid ambiguity.

Where:

- *b*_obs_ denotes the observed barcode sequence from sequencing output, potentially containing sequencing errors.
- *b*_*p*_ refers to a barcode in the empirically derived permit list *P*.
- *P* represents the set of trusted barcodes identified from the current dataset using the defined threshold θ.
- count(*b*) indicates the total UMI count observed for barcode *b*.
- *θ* specifies the UMI count threshold above which barcodes are considered valid and included in *P*.
- Distance criterion defines the maximum allowable mismatches (e.g., Hamming distance ≤□1) for correction to occur.

Symbolically: b? → [Histogram | θ] → Permit List → b⍰

Where:

- *b*? denotes the barcode of uncertain accuracy.
- Histogram | *θ* indicates the data-driven thresholding process used to generate the permit list from the barcode count distribution.
- Permit List represents the set *P* of barcodes included after thresholding.
- *b*⍰ denotes the corrected barcode assigned from exactly one matching entry in *P* within the distance threshold.

### UMI Deduplication

The final core stage of primary processing is Unique Molecular Identifier (UMI) deduplication, which corrects for PCR amplification bias by ensuring that multiple reads originating from a single RNA molecule are counted only once (Fig. 1d). This step is performed on a per-cell, per-gene basis. The benchmarked pipelines employ four distinct strategies for handling potential sequencing errors within UMI sequences, ranging from active error correction to conservative exact matching.

The deduplication rules for a set of observed UMIs U_obs within a given cell-gene pair can be formalized as follows:

#### 1. Cell Ranger (Directional Graph-based Correction)

To correct sequencing errors in Unique Molecular Identifiers (UMIs) while minimizing over-merging, Cell Ranger uses a directional, abundance-aware graph-based algorithm. All observed UMIs for a given cell barcode and gene are represented as nodes in a UMI graph. An edge is drawn between any two nodes if their Hamming distance *H* is 1 (i.e., they differ by exactly one nucleotide).

After building the graph, Cell Ranger partitions it into connected components *C*, where each component contains UMIs potentially representing the same original molecule. Within each connected component, UMIs are merged in a directional fashion from lower-abundance nodes toward higher-abundance nodes.

This is achieved by selecting the most abundant UMI in the component as the representative UMI *u*_rep_:

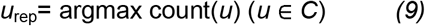

Here, count(*u*) is the number of reads associated with UMI *u*. All other lower-abundance UMIs *u*_low_ in *C* are collapsed into *u*_rep_ if they satisfy the directional abundance condition-typically that count(*u*_high_) is at least double that of count(*u*_low_), ensuring that errors are corrected toward genuinely abundant UMIs.

This directional merging corrects low-abundance UMIs likely arising from sequencing errors while preserving distinct molecules in cases of similar abundance.

Where:

- *u*_low_ denotes a UMI node in the graph that has a lower read count and is a candidate for correction.
- *u*_high_ denotes a UMI node in the graph that has a higher read count and serves as the potential error-free target for merging.
- *H*≤1 specifies that the Hamming distance threshold allows an edge only if two UMIs differ by at most one base.
- Graph represents the network structure where the nodes correspond to UMIs and the edges represent potential single-base sequencing errors.
- Connected component *C* defines the subset of the graph in which every two UMIs are connected by a path composed of Hamming-distance-1 edges.
- count(*u*) indicates the number of sequencing reads associated with UMI *u*.
- argmax returns the UMI within *C* that has the maximum read count.
- *u*_rep_ identifies the representative UMI for the component, into which all UMIs in *C* are collapsed if the abundance rule is satisfied.

Symbolically: UMI_low_ → [H≤1 | Graph-Abundance] → UMI_high_

Where:

- UMI_low_ refers to the observed UMI with fewer supporting reads that is considered a candidate for correction or merging.
- *H*≤1 specifies that the Hamming distance between the sequences of UMI_low_ and UMI_high_ must be one or zero (i.e., they differ by at most one nucleotide).
- Graph-Abundance indicates that the correction decision is based on a graph-based traversal of UMIs connected by Hamming-distance-1 edges, where abundance (read count) guides the merging direction.
- UMI_high_ refers to the UMI with higher read count within the connected component, representing the presumed true (error-free) UMI into which UMI_low_ is merged.

#### 2. STARsolo (Directional Heuristic-based Correction)

STARsolo employs a conceptually similar directional, abundance-aware strategy but relies on a simpler and more conservative heuristic instead of a full graph traversal. This method avoids some complex merging decisions that can occur in dense graph components. The default directional mode merges a less abundant UMI into a more abundant one if they are within a Hamming distance of 1 and the more abundant UMI is supported by at least twice as many reads. (Default directional mode) For any two UMIs *u*_*i*_, *u*_*j*_ ∈ *U*_obs_, merge *u*_*j*_ into *u*_*i*_ if:

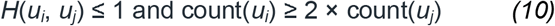

Where:

- *U*_obs_ refers to the set of all UMIs observed for a specific cell barcode and gene.
- *u*_*i*_ denotes the more abundant UMI candidate for retaining after correction.
- *u*_*j*_ denotes the less abundant UMI candidate considered for merging into *u*_*i*_.
- *H*(*u*_*i*_, *u*_*j*_) represents the Hamming distance between *u*_*i*_ and *u*_*j*_, defined as the number of nucleotide positions at which the sequences differ.
- ≤1 specifies the maximum Hamming distance allowed for considering a merge, meaning one or fewer mismatches.
- count(*u*) indicates the number of sequencing reads associated with the UMI *u*.
- ≥2× describes the abundance threshold that the more abundant UMI must exceed in relation to the less abundant UMI for a merge to occur.

Symbolically: UMI_low_ → [H≤1 | Heuristic-Abundance] → UMI_high_

Where:

- UMI_low_ refers to the observed UMI with fewer reads, considered an error candidate.
- *H*≤1 specifies that the UMI sequences must differ by no more than one base to be eligible for merging.
- Heuristic-Abundance indicates that merging is governed by a simple abundance ratio requirement rather than full graph analysis.
- UMI_high_ refers to the corrected UMI with higher abundance that serves as the merge target.

#### 3. KBus (Exact Match)

KBus applies a conservative, count-agnostic strategy for Unique Molecular Identifier (UMI) deduplication that avoids any sequence-based error correction. Under this method, UMIs are collapsed only if they are identical in sequence.

For each cell barcode-gene pair, KBus groups all reads that share the same cell barcode, same UMI sequence, and same gene assignment, collapsing them into a single molecule count.

Sequence similarity measures (e.g., Hamming distance) and abundance ratios play no role in this process.

The final molecular count for each cell-gene pair is simply the number of distinct UMI sequences present:

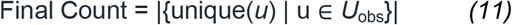

Symbolically: UMI → [Exact Match] → Count

Where:

- *U*_obs_ refers to the set of all UMIs observed for a given cell barcode and gene.
- unique(*u*) identifies a UMI sequence that is distinct from all other UMIs in the same *U*_obs_ set.
- ∣⋅∣ denotes the number of elements in a set.
- Count refers to the total number of distinct UMIs, representing molecular counts.

#### 4. Alevin-fry (Exact Match)

In its default and most commonly used modes, Alevin-fry employs the same exact match approach used by KBus for UMI deduplication. Under this policy, no explicit sequence-based error correction is performed; every unique UMI sequence represents a distinct molecule.

For each cell barcode–gene combination, Alevin-fry groups reads that share identical cell barcodes, identical UMI sequences, and identical gene annotation, collapsing each group into a single molecule count. Unlike error-correcting strategies, this method leaves potentially erroneous UMIs unmerged if they differ by even a single nucleotide.

The molecule count is calculated using the same rule as KBus:

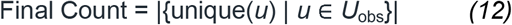

Symbolically: UMI → [Exact Match] → Count Where:

- *U*_obs_ refers to the set of all UMIs observed for a given cell barcode and gene.
- unique(*u*) identifies a UMI sequence that is distinct from all other UMIs in the same *U*_obs_ set.
- ∣⋅∣ denotes the number of elements in a set.
- Count refers to the total number of distinct UMIs, representing molecular counts.

### CAS Calculation

To quantitatively evaluate the concordance between direct cell-level annotation and cluster-level consensus annotation, we developed the Cluster Agreement Score (CAS). CAS measures the proportion of cells in each cluster whose direct annotation agrees with the consensus label assigned to that cluster.

For each Leiden cluster *C*, we:

1. Determine its consensus label *L*_*C*_, defined as the majority label among all direct annotations of cells in *C*.
2. For each cell *j* ∈ *C*, compare the cell’s direct annotation *l*_*j*_ to *L*_*C*_.
3. Compute the fraction of cells in *C* where *l*_*j*_ = *L*_*C*_:

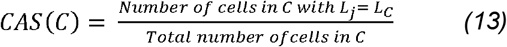

We then compute the overall CAS across all *K* clusters as:

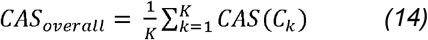

Where:

- *C*_*k*_ = the *k*-th cluster.
- *L*_*Ck*_ = consensus annotation label for cluster *C*_*k*_.
- *l*_*j*_ = direct annotation for cell *j*.
- *K* = total number of clusters.

The resulting CAS ranges from 0 to 1:

- 1 indicates perfect agreement-all cells in all clusters share the same label as their cluster’s consensus.
- 0 indicates complete disagreement -no cell’s direct annotation matches its cluster’s consensus label.

### MCS Calculation

To quantitatively assess annotation accuracy at the gene level, we developed the MCS, which measures the degree to which an annotated cell type exhibits its expected canonical gene expression signature in the post-processing data. For each annotated cell type *T*, we identified its corresponding cluster *C* and retrieved the complete set of *n* canonical marker genes{*m*_1_, *m*_2_, … , *m*_*n*_} defined for that type. Expression prevalence for a marker *m*_*i*_ was calculated as the fraction of cells in *C* with non-zero log-normalized expression for *m*_*i*_. The MCS for *T* was then computed as:

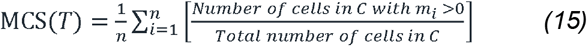

Where:

- *T* = annotated cell type.
- *C* = cluster assigned to *T*.
- *m*_*i*_ = *i*-th canonical marker gene for *T*.
- *n* = number of canonical marker genes for *T*.
- Expression *m*_*i*_>0 refers to non-zero log-normalized expression counts.

The resulting MCS value ranges from 0 to 1, where 1 indicates that all cells in *C* express all canonical markers for *T*, and 0 indicates complete absence of marker expression.

### Data analysis

#### 1. Data processing. For snRNA-seq analysis, processed reads were aligned using STAR

(v2.7.11b), kallisto (v0.51.0), or Cell Ranger (v9.0.1), and downstream analyses were conducted with Scanpy (v1.11.1). Simulated datasets were generated with Splatter (v1.26.0) and polyester (v1.38.0). All steps used reference genome resources from the Ensembl Mus musculus GRCm39 assembly, including Mus_musculus.GRCm39.dna.primary.assembly.fa, Mus_musculus.GRCm39.cdna.all.fa, and Mus_musculus.GRCm39.112.chr.gtf.

#### 2. scRNA-seq Data Quantification and Cell-Gene Matrix Generation

Raw single-cell RNA sequencing reads were processed and quantified to generate cell-gene count matrices using three different approaches:

##### 2.1 Cell Ranger

Single-cell RNA sequencing reads were processed and quantified using the 10x Genomics Cell Ranger pipeline. The cellranger count command was executed to align reads to the reference transcriptome, perform cell identification, and generate the corresponding cell-gene expression matrix. The specific command used was:

cellranger count --id=5k_mouse_brain_CNIK_3pv3_output -- sample=5k_mouse_brain_CNIK_3pv3 --fastqs=/home/data/qs/mus_str.sn/HGC20231021002-0003-2/cellranger_output/ --transcriptome=/home/data/qs/tools/refdata-gex-GRCm39-2024-A/ -- localcores=40 --localmem=200 --expect-cells=15000 --create-bam true

##### 2.2 STARsolo

Single-cell sequencing reads were aligned and quantified using STAR in STARsolo mode. In this workflow, reads were processed to detect cell barcodes and unique molecular identifiers (UMIs), aligned to the reference genome index, and quantified for gene expression based on UMI counts. The command parameters were configured for 10x Genomics chemistry, specifying cell barcode and UMI positions and lengths, forward-strand specificity, UMI deduplication with a tolerance of one mismatch, and the use of a provided cell barcode whitelist. The full command executed was:

STAR --runThreadN 40 --runMode alignReads --soloType CB_UMI_Simple --soloCBstart 1 -- soloCBlen 16 --soloUMIstart 17 --soloUMIlen 12 --soloBarcodeReadLength 0 --readFilesIn 5k_mouse_brain_CNIK_3pv3_S2_L001_R2_001.fastq.gz 5k_mouse_brain_CNIK_3pv3_S2_L001_R1_001.fastq.gz --readFilesCommand zcat –genomeDir /home/data/qs/data/reference_isoform/mus_ref/STAR_index/ --outFileNamePrefix /path/STARsolo/musWT_STR_ --outSAMtype BAM SortedByCoordinate --soloFeatures GeneFull_Ex50pAS --soloUMIdedup 1MM_CR --soloStrand Forward --soloCBmatchWLtype 1MM_multi_Nbase_pseudocounts --soloCBwhitelist /path/3M-february-2018.txt

##### 2.3 KBus Workflow

The KBus workflow was utilized for quantification. The steps were executed sequentially:

###### 2.3.1 The analysis was performed within a dedicated conda environment

conda activate kb_env

###### 2.3.2 A kallisto index was built from the reference cDNA sequence

kallisto index -i /path/mus_ref/Mus_musculus.GRCm39_kallito.idx /path/mus_ref/Mus_musculus.GRCm39.cdna.all.fa.gz

###### 2.3.3 Reads were pseudoaligned to the index and converted into a BUS file

kallisto bus -i /path/mus_ref/Mus_musculus.GRCm39_kallito.idx -o ./kbus_output -x 10xv3 -t 8 5k_mouse_brain_CNIK_3pv3_S2_L001_R1_001.fastq.gz 5k_mouse_brain_CNIK_3pv3_S2_L001_R2_001.fastq.gz

This command specified the kallisto index (-i), output directory (-o), single-cell chemistry (-x 10xv3), number of threads (-t), and the input FASTQ files.

###### 2.3.4 Cell barcodes in the BUS file were corrected against a provided whitelist

bustools correct -w /path/3M-february-2018.txt -o /path/kbus_output/corrected.bus /path/kbus_output/output.bus

###### 2.3.5 The corrected BUS file was sorted

bustools sort -o /path/kbus_output/sorted.bus /path/kbus_output/corrected.bus

###### 2.3.6 Gene-level UMI counts were generated per cell

bustools count -o /path/kbus_output/counts -g /path/tx2gene_with_version.tsv -e /path/kbus_output/matrix.ec -t /path/kbus_output/transcripts.txt –genecounts /path/kbus_output/sorted.bus

This final step used the sorted BUS file, providing the transcript-to-gene mapping (-g), equivalence classes (-e), and transcript list (-t) to aggregate counts to the gene level (-- genecounts).

##### 2.4 Salmon and Alevin-fry Workflow

###### 2.4.1 Alignment and RAD File Generation with Alevin-fry

salmon alevin -l ISR -i “/home/data/qs/mus_str.sn/HGC20231021002-0003-1/salmon_index/Mus_musculus.GRCm39.cdna.all_salmon_index_unversioned_v2/” -1 “./5k_mouse_brain_CNIK_3pv3_S2_L001_R1_001.fastq.gz” -2 “./5k_mouse_brain_CNIK_3pv3_S2_L001_R2_001.fastq.gz” -p 40 -o “./result_Alevin/5k_mouse_brain_CNIK_3pv3_alevin_rad_output/” --tgMap “/home/data/qs/mus_str.sn/HGC20231021002-0003-1/salmon_alevin_index/txp2gene.tsv” -- chromiumV3 --rad

###### 2.4.2 Permit List Generation

Alevin-fry was used to generate a whitelist of valid cell barcodes:

alevin-fry generate-permit-list -d fw -i “./5k_mouse_brain_CNIK_3pv3_alevin_rad_output/” -o “./5k_mouse_brain_CNIK_3pv3_alevin_fry_output/” -k

###### 2.4.3 Alternative Permit List Generation (Direct Output)

An alternative run outputting the permit list directly into the RAD output folder:

alevin-fry generate-permit-list -d fw -i “./5k_mouse_brain_CNIK_3pv3_alevin_rad_output/” -o “./5k_mouse_brain_CNIK_3pv3_alevin_rad_output/” -k

###### 2.4.4 Copying Intermediate Files

Intermediate files from the RAD output directory were copied into the fry output directory for subsequent steps:

cp ./5k_mouse_brain_CNIK_3pv3_alevin_rad_output/^*^ ./5k_mouse_brain_CNIK_3pv3_alevin_fry_output/

###### 2.4.5 Collation Step

Collation was performed to reorganize RAD file data:

alevin-fry collate -i “./5k_mouse_brain_CNIK_3pv3_alevin_rad_output/” -r “./5k_mouse_brain_CNIK_3pv3_alevin_fry_output/” -t 40

#### 3. Generation of Simulated scRNA-seq Data

To provide a ground-truth dataset for validation, a two-step pipeline was developed to generate Cell Ranger-compatible simulated scRNA-seq data. The first step, executed by the R script step1_simulate_counts_fastq.R, uses the splatter package to simulate a count matrix and the polyester package to generate corresponding raw FASTQ reads. The second step, handled by the Python script step2_add_barcodes.py, processes these reads by adding authentic 10x Genomics cell barcodes and UMIs, formatting them into the R1/R2 file structure required by Cell Ranger. Both command-line scripts are publicly available in the simulation/directory of the project’s GitHub repository: https://github.com/QiangSu/CAS-MCS-Scoring.

#### 4. Automated Cell Type Annotation using Scanpy and CellTypist

Post-quantification analysis was conducted using a suite of custom Python scripts built upon the Scanpy library (v1.9.6). These modular scripts replace an initial exploratory Jupyter notebook analysis and provide reproducible, command-line workflows. The repository includes the following pipelines:

- Annotation Quality Scoring (CAS-MCS-Scoring.py): This streamlined utility is ideal for quickly assessing annotation confidence.
- Post-Analysis Purity Assessment (calculate_cluster_purity.py): A utility script for quality control, designed to calculate the annotation consistency for each cluster using the output from the per-cell annotation pipeline.
- Single-Sample Consensus Annotation (scanpy_pipeline_majority_voting.py): This pipeline processes a single dataset, performing clustering and then assigning a single, consensus cell type label to all cells within each cluster based on a “majority vote” of CellTypist predictions.
- Single-Sample Per-Cell Annotation (scanpy_pipeline_per_cell.py): This parallel pipeline also processes a single dataset, but assigns an independent CellTypist label to each individual cell, allowing for the investigation of intra-cluster heterogeneity.
- Multi-Sample Integration and DGE (run_integration_analysis.py): This advanced workflow is designed for comparative analysis of two or more samples. It uses Harmony to correct for batch effects before performing joint dimensionality reduction, clustering, and cell type annotation. Crucially, it also conducts differential gene expression (DGE) analysis between user-defined conditions within each identified cell type.

Comprehensive single-cell analysis was performed using the Scanpy Python library. The entire methodology was implemented as a suite of modular Python scripts, encompassing standard preprocessing of cell-gene matrices, graph-based clustering using the Leiden algorithm, and advanced dimensionality reduction techniques like UMAP for visualizing distinct cell populations. Automated cell type annotation was performed using CellTypist. The complete analysis workflow, including scripts for multi-sample integration and differential gene expression, is publicly accessible within this repository (https://github.com/QiangSu/CAS-MCS-Scoring).

##### 4.1 Data Loading and Quality Control (Both Pipelines)

Raw 10x Genomics gene expression matrices (matrix.mtx.gz, barcodes.tsv.gz, features.tsv.gz) were loaded using scanpy.read_10x_mtx with var_names=‘gene_symbols’. Gene symbols were made unique and raw count data stored in adata.layers[“counts”]. Quality control (QC) metrics were computed with scanpy.pp.calculate_qc_metrics, marking mitochondrial genes using the prefix mt-. QC visualizations (violin plots for gene counts, total counts, mitochondrial percentage; scatter plots for total counts vs detected genes) were generated prior to filtering. Cells were retained if they contained 200–7000 detected genes and <10% of counts from mitochondrial reads. Genes expressed in fewer than 3 cells were excluded.

##### 4.2 Normalization, Feature Selection, and Dimensionality Reduction (Both Pipelines)

Filtered counts were normalized to 10,000 counts per cell (scanpy.pp.normalize_total) and log-transformed (scanpy.pp.log1p). Highly variable genes (HVGs) were identified using the Seurat v3 method (scanpy.pp.highly_variable_genes, n_top_genes=10000) and retained for downstream analysis, while the full normalized dataset was stored in adata.raw. The HVG matrix was scaled (scanpy.pp.scale, max_value=10) and subjected to Principal Component Analysis (scanpy.tl.pca, 50 components). PCA variance ratios were plotted to assess dimensionality.

##### 4.3 Clustering and Cell Type Annotation

###### 4.3.1 Graph-Based Clustering

For both pipelines, a neighborhood graph (scanpy.pp.neighbors) was constructed using the top 30 PCs and 15 nearest neighbors. Leiden clustering (scanpy.tl.leiden, resolution□=□0.9) was applied to partition the cells. UMAP embeddings (scanpy.tl.umap) were computed for visualization, and silhouette scores were calculated from PCA space to evaluate cluster compactness.

###### 4.3.2 CellTypist Annotation-Pipeline 1 (Majority Voting Enabled)

In scanpy_pipeline.py, the Mouse_Whole_Brain.pkl CellTypist model was loaded, and cell-type predictions were obtained via celltypist.annotate with majority_voting=True and mode=‘best match’. Each Leiden cluster was assigned a consensus cell-type label using majority vote across its member cells

##### 4.4 Data Export

The results of each pipeline were comprehensively exported. This included the raw per-cell annotation tables from CellTypist and curated consensus annotation tables in CSV format. The final, fully annotated AnnData object for each analysis, containing all metadata, dimensionality reduction embeddings, and clustering results, was saved in the .h5ad file format.

### Statistics

Statistical analyses were performed as appropriate for each dataset. Linear relationships were assessed using Pearson correlation coefficients and adjusted R^2^ values to evaluate model fit. Marker gene identification in the sc/snRNA-seq data was carried out using the Wilcoxon rank-sum test (Mann–Whitney U test), comparing the expression of each gene in a given annotated cell type against all other cells to rank candidate markers. Multiple testing corrections were applied to the resulting p-values, with significance defined as adjusted p□<□0.05.

### Data access

The raw sequencing data have been deposited in the NCBI Sequence Read Archive (SRA; https://www.ncbi.nlm.nih.gov/sra) under project accession PRJNA1263897. The snRNA-seq datasets for musHuntingtonD-STR and musWT-STR are available under this project with SRA accessions SRR33607933 and SRR33607934, respectively. Additional publicly available datasets, identified by their corresponding SRR accessions, were obtained from NCBI. While some of these accessions are cited in their original publications, others correspond to ongoing, unpublished studies and are not yet formally referenced.

## Software availability

All code developed for this study is open-source and publicly available in the CAS-MCS-Scoring GitHub repository (https://github.com/QiangSu/CAS-MCS-Scoring). The primary contribution of this repository is the CAS-MCS-Scoring.py utility, a reusable command-line pipeline for the quantitative evaluation of cell type annotation accuracy. Located in the scripts/directory, this tool implements the dual-metric framework central to our study’s benchmarking approach: the CAS for measuring internal annotation consistency and the MCS for assessing external biological validation. The repository also includes a comprehensive suite of supporting tools. The scripts/directory contains modular Python pipelines for standard single-cell workflows, such as quality control, two-sample integration, and differential gene expression analysis. A separate R and Python-based workflow for the de novo simulation of Cell Ranger-compatible FASTQ files is provided in the simulation/directory. Reproducibility of the computational environment is ensured by a Conda environment file (environment/environment.yml). All scripts are implemented as configurable command-line tools to facilitate ease of use and adaptation.

## Acknowledgments

We acknowledge financial support from the National Key Research and Development Program of China (2023YFA0914904, 2022YFA1105601).

## Author Contributions

Qiang Su conceived and designed the study, performed all data analyses, and generated the figures. Qiang Su and Yi Long drafted the manuscript, with input and critical revisions from all authors. Fuyu Duan provided raw data. Qiang Su and Qizhou Lian supervised the project. All authors reviewed and approved the final version of the manuscript.

